# Low RT-based Genome Editing Fidelity in Mouse Hepatocytes: Challenges and Solutions

**DOI:** 10.1101/2024.10.31.621282

**Authors:** Maike Thamsen Dunyak, Patrick Hanna, Angela X. Nan, Jessica von Stetina, Rejina Pokharel, Mollie O’Hara, Brett J.G. Estes, Kangni Zheng, Sonke Svenson, Jason Andresen, Wei Li, Mini Agrawal, Chase Hughes, Makayla Spicer, Vimal Choudhary, Lily Kelley, Jie Wang, Martin Meagher, Jenny Xie, Swati Mukherjee, Jonathan D. Finn

## Abstract

Integrase-mediated Programmable Genomic Integration (I-PGI) uses a Cas9 nickase (nCas9) with a reverse transcriptase (RT), to write a large serine integrase (LSI) target site (attB/P, here called “beacon”) in a programmed location. Co-delivery of the LSI and a DNA template containing the cognate recognition site results in precise integration of the template in a specific genomic location. While we were able to achieve high-fidelity beacon placement in a range of primate cycling and non-dividing cells, when translating our technology into an *in vivo* rodent model (liver) we surprisingly observed very low beacon fidelity, with the vast majority of beacons being unsuitable for integration. This phenomenon was independent of mouse strain, but was specific to non-dividing cells, as a cycling mouse hepatocyte cell line (Hepa1-6) demonstrated very high levels of fidelity. To address this issue we utilized neonatal mice, which have a much higher proportion of proliferating hepatocytes than adult mice. This resulted in a significant increase in the placement of high-fidelity beacons, and achieved functional gene expression after I-PGI in a therapeutically relevant target site. In an alternate approach, we engineered transgenic mice with intact beacons placed in specific genomic locations, allowing us to optimize integrase and DNA template dosing and kinetics. In summary, we have identified a previously undescribed challenge when using RT-based editing to write long sequences (~40 bp) in non-dividing rodent hepatocytes. This phenomenon was specific to rodents and was not observed in primate dividing or non-dividing cells. This previously unidentified challenge using RTs will limit the use of I-PGI in mouse models, however here we describe two methods that address this issue.

## Introduction

Over the past decade, genome editing has had a major impact across many fields, including basic and translational research [1–3]. What is even more remarkable is that this incredible progress has been made with fairly limited gene editing tools, generally limited to either breaking genes (via error-prone repair of double-stranded DNA breaks (DSBs), or making very small edits (1-3 nucleotide (nt)) to either knock-out genes/regulatory elements or correct very specific mutations [4, 5]. In order to truly harness the power of gene editing, we must go beyond these small edits and move towards programmable genomic integration (PGI), which is the ability to efficiently insert large pieces of DNA into specific genomic locations. This capability would enable mutation agnostic approaches to cure genetic diseases by inserting healthy copies of genes in the correct genomic location, and unleash the field of cell engineering, allowing the precise integration of logic gates, safety switches, or novel regulatory elements.

Our team has focused on the development of a recent innovation that combines two previous editing technologies to enable the precise integration of large DNA sequences. The first description of this, PASTE (Programmable Addition via Site-specific Targeting Elements) combines the programmability of a Cas9 nickase fused with a reverse transcriptase (RT) enzyme to first write a short sequence (~40 bp, here called a ‘beacon’) at a user defined location in the genome [6] (Fig 1A). This sequence (attP or attB) is the recognition site for a specific large serine integrase (LSI). When the LSI is delivered in the presence of dsDNA that contains the cognate recognition sequence (attB or attP), the LSI is able to integrate the template DNA sequence at the target site in the genome. Importantly, the integrase-based insertion approach is agnostic to the size of the DNA insert, meaning that the only restriction on the size of the DNA insert is delivery efficiency. While this is a simple concept, the clever use of three enzymes (nCas9, RT, LSI) in a novel way allowed for an elegant solution to a problem that has plagued the gene therapy field for decades: how to efficiently put a large piece of DNA in a single specific genomic location.

**Fig 1.**
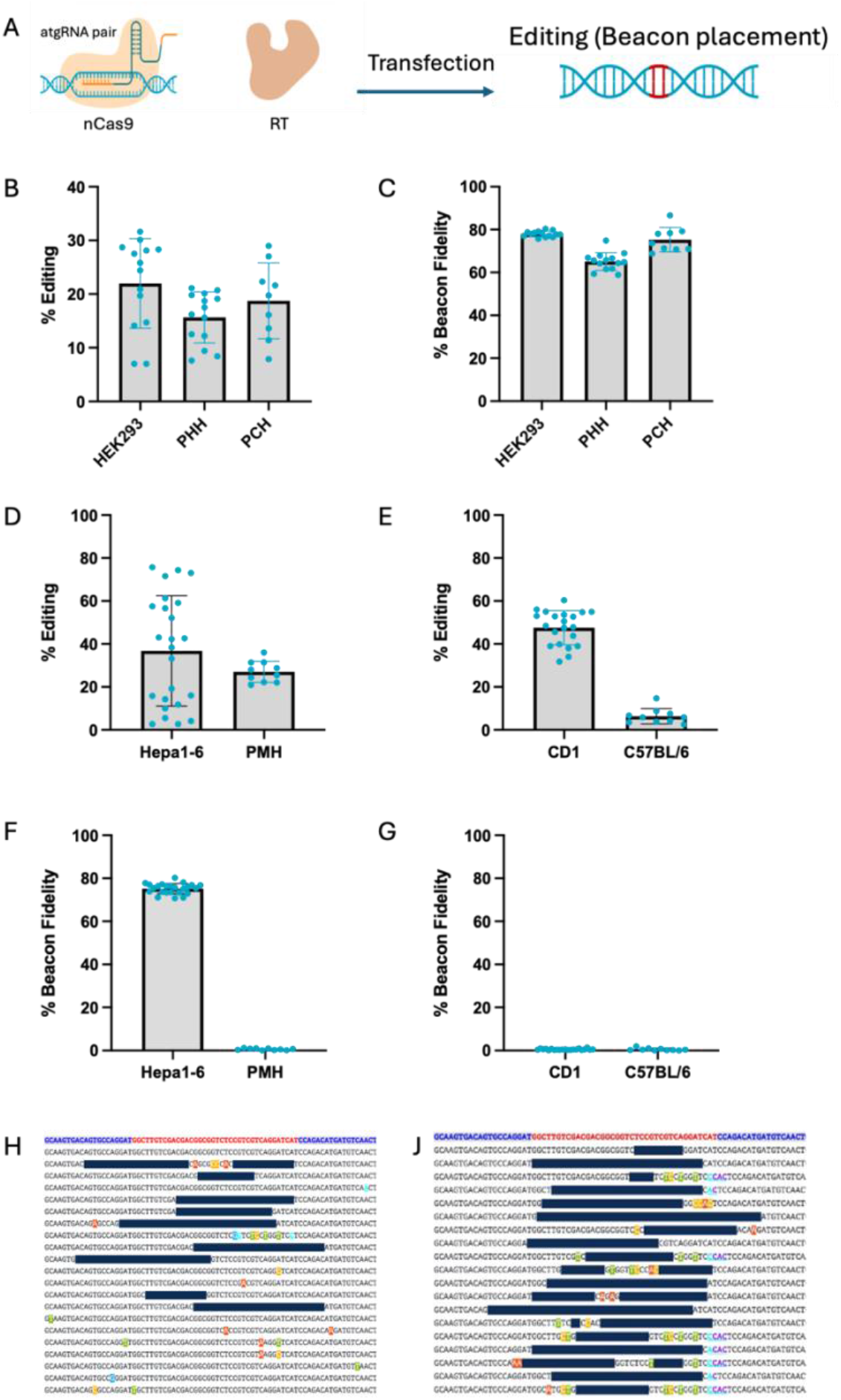
In vitro and in vivo editing potency and editing fidelity for beacon placement. Diagram showing beacon placement editing with LNPs or transfection (A) In vitro editing potency (B) and fidelity (C) in human and cynomolgus cells. In vitro (D) and in vivo editing potency (E) and fidelity (F&G) in murine cells. Cells were transfected with Messenger Max (MM) and mice treated IV with LNPs (3mg/kg). LNPs contained nCas9-RT mRNA and atgRNAs. Multiple sequence alignment of editing in Hepa1-6 cells (F) and CD1 mouse (G).

This approach has shown promise *in vitro* across a range of cycling cancer cell lines [6, 7], and we have recently described the translation of this system to non-human primates [8]. Surprisingly when we were translating this system to mouse models, we uncovered some unexpected challenges that severely limited the utility of this approach. These challenges, and potential solutions, are described here.

## Results and Discussion

While previous groups have demonstrated that RT-based editing systems can efficiently make small edits in a variety of cell types, [9, 10] to date no one has demonstrated efficient writing of large (>10 bp) sequences in non-dividing primate or rodent cells. After significant optimization through protein engineering and guide architecture/chemical modification (data not shown), we were able to efficiently write beacons (~40 bp) in cycling (HEK293) and non-cycling (primary human hepatocytes (PHH) or primary cynomolgus hepatocytes (PCH)) primate cells (Fig 1B). The majority of beacons written had excellent fidelity, with >70% of ed being the correct LSI recognition sequence (Fig 1C). Using lipid nanoparticles (LNPs) to deliver the beacon writing machinery (nCas9-RT mRNA + dual attachment guide RNA (atgRNAs) [8], we were able to achieve efficient beacon placement (~40%) in mouse liver following a single intravenous (i.v.) administration (Fig 1E). However, we found that the fidelity of the beacons was very low, with less than 1% of the beacons being intact and able to be recognized by the LSI (Fig 1G, J). The poor beacon fidelity was independent of the mouse strain used and was also observed *in vitro* in non-dividing primary mouse hepatocytes (PMH) (Fig 1D, F, H). In general, we found that the beacons in PMH were either truncated (incorporated half of the beacon sequence) or consisted of excisions between the paired nick sites. Given the discordance in fidelity between the primate and mouse hepatocytes, we decided to investigate further. We found that this was not simply due to a difference in species, as when using Hepa1-6 cells (cycling mouse hepatocyte hepatoma cell line) we observed high levels of beacon fidelity (>70%) (Fig 1F, H).

Based on these results, we speculated that the challenges with beacon fidelity could be related to the non-dividing nature of primary mouse hepatocytes. There are many differences between dividing and non-dividing cells, including differences in DNA repair pathways (e.g. HDR vs NHEJ) [11] and intracellular dNTP pools [12, 13]. Given the reliance of RT-based editing on both host DNA repair machinery and the concentration of dNTPs (used by the RT for beacon writing), it is possible that multiple mechanisms may be responsible for the poor beacon fidelity in mouse hepatocytes. To test this hypothesis, we treated neonatal mice (Day 2 postnatal), as a significant number of hepatocytes are undergoing cell division during the first days of life [14]. We were able to achieve comparable levels of beacon placement in neonates and adult mice, however we observed a significant (40-fold) increase in beacon fidelity in neonates, resulting in up to 48% fidelity and 33% intact beacons in bulk liver (Fig 2A). While this level of fidelity is lower than in Hepa1-6 cells (Fig 1F), the dramatic increase in fidelity when treating neonates supports the hypothesis that the poor fidelity in adult mice is related to the terminally differentiated non-dividing nature of adult hepatocytes.

**Fig 2.**
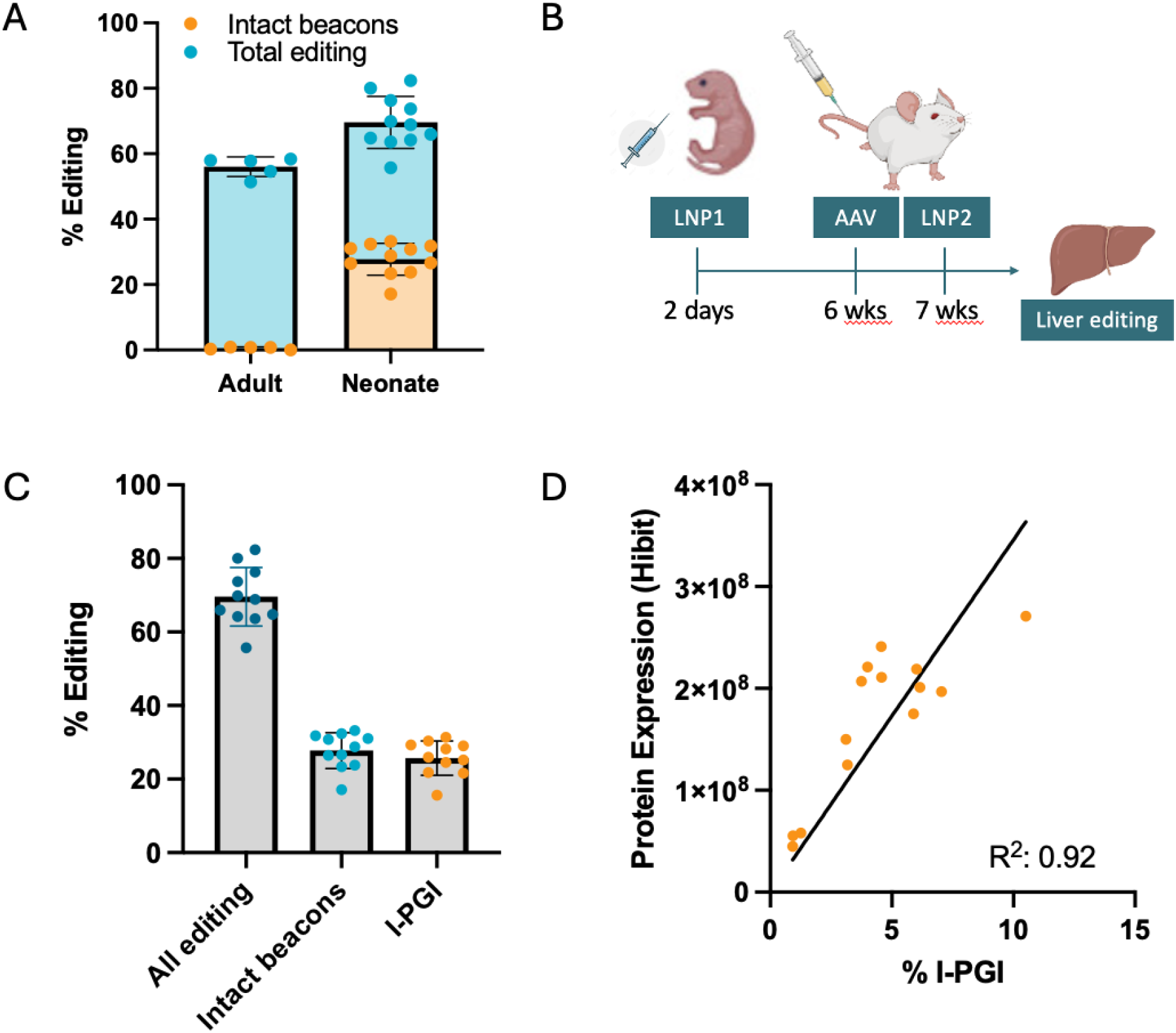
Beacon fidelity and editing (I-PGI) in neonatal mice. Comparison of editing fidelity in adult and neonatal mice treated with 3mk/kg LNPs containing nCas9-RT mRNA and atgRNAs (LNP1) to facilitate beacon placement in mouse Pah (A). Diagram outlining dosing regimen for neonatal I-PGI studies (B). Beacon placement was performed using LNPs containing nCas9-RT mRNA and atgRNAs (LNP1, 3-6mg/kg) by IV injection into neonates postnatal day 2. AAV template (2e13-5e13vg/kg) was administered when mice reached 6 weeks of age followed by Integrase mRNA LNP (LNP2, 3-6mg/kg) at 7 weeks. Analysis was performed on liver tissue and serum at week 8. C shows I-PGI in neonatal mice integrating human Pah in the mouse Pah locus. ‘All editing’ includes Indels, partial beacons, intact beacons (assessed by NGS) and integration events (assessed by ddPCR). ‘Intact beacons’ shows the percent of alleles that had intact beacons and I-PGI shows percent of alleles with successful integration. Detection of Hibit-tagged protein in serum of mice after I-PGI. A Hibit-tagged human Factor 9 template was integrated in the mouse Pah locus.

Given that we were now able to achieve reasonable levels of intact beacons in mice, we were able to use a combination of LNP delivery of LSI mRNA and AAV delivery of template DNA to achieve full PGI *in vivo* (Fig 2B). We targeted intron 1 of the mouse phenylalanine hydroxylase (PAH) gene and inserted a HiBiT-tagged human Factor 9 (hF9-HiBiT) reporter template containing a splice acceptor that would result in expression of the inserted reporter gene regulated by the endogenous PAH promoter (Fig 2C). As expected, we observed a strong correlation (R^2^=0.92) between I-PGI levels and reporter gene expression (Fig 2D).

While writing beacons in neonatal mice was able to achieve functional PGI, the challenges associated with neonatal dosing and the long duration for each experiment (eight weeks) led us to explore alternate methods for LSI-mediated gene integration in mice. To overcome the beacon fidelity issue, we generated transgenic mouse models that had pre-installed beacons in therapeutically relevant sites (Rosa26, PAH, F9). Using F9 as a proof of concept (PoC) gene, we were able to achieve efficient levels (27%) of PGI using LNP delivery of LSI mRNA and AAV delivery of an hF9 template (exon 2-8, including a splice acceptor and polyA, Fig 3A-C). Integration of hF9 into the murine F9 intron 1 resulted in transcription, splicing, translation, and post-translational processing of the integrated transgene, achieving serum hF9 levels averaging 50% of normal human levels (Fig 3D). Using an antibody that can discriminate between human and murine F9, we performed immunohistochemistry (IHC) on liver sections from treated mice and observed robust staining of hF9 throughout the mouse liver (Fig 3E-G). Interestingly, the pattern of hF9 staining in treated mice closely resembled the pattern of hF9 staining from a normal human liver, with the majority of signal coming from the sinusoidal space. These data suggest that expressing hF9 from the native promoter was able to maintain endogenous regulation and protein expression levels. When we quantified the number of hF9 positive cells (area-based analysis), we were able to achieve expression in ~45-50% of cells, which correlated with the serum levels of hF9 (Fig 3G). A possible explanation for the discrepancy between hF9 positive cells and PGI levels could be monoallelic editing of cells as the majority of hepatocytes in adult mice are polyploid [15], and thus editing of a single allele will result in a hF9 positive cell. Integrase-mediated PGI was not restricted to the F9 gene, as efficient integration was also observed in two additional loci (Rosa26, PAH) (Fig S1).

**Fig 3.**
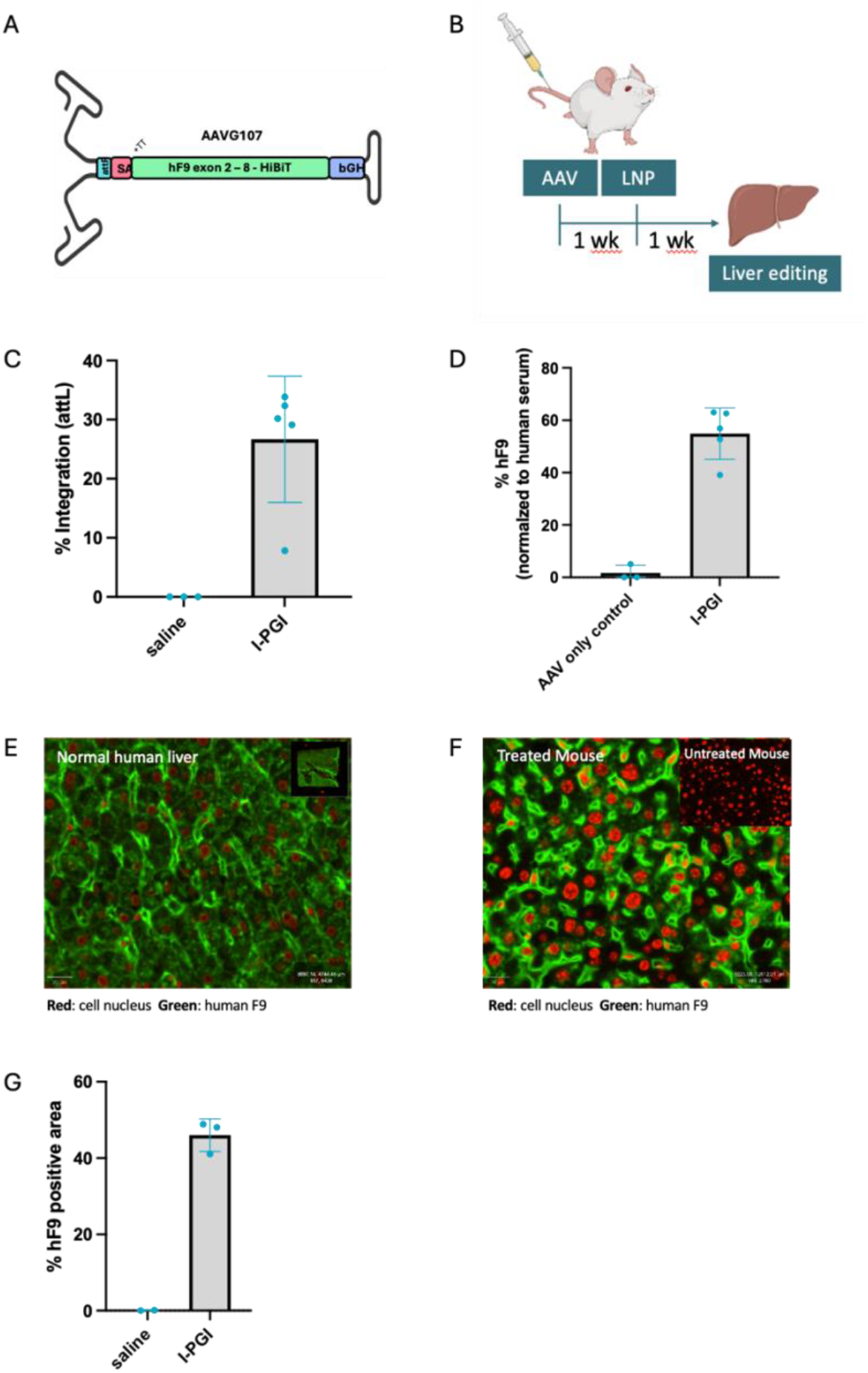
I-PGI in transgenic attB beacon mice. AAV DNA template encoding fF9 (A). Mice were treated with AAV containing the template DNA (2e13-2.5e13vg/kg) and 1 week later with 3mg/kg LNPs containing Integrase mRNA (B). Liver and serum were collected 1 week post LNP dosing. Integration efficiency in male transgenic mice with intact attB site placed in Intron 1 of Factor 9. Integration measured by ddPCR and normalized to a mouse F9 reference (C). Human factor 9 levels measured by ELISA in serum of Factor 9 attB mice after I-PGI (D). Levels are normalized to Factor 9 levels in human serum. Immunofluorescent imaging of human Factor 9 in normal human liver (E) and in liver of a treated Factor 9 attB mouse after I-PGI (E). Image of human factor 9 staining in an untreated Factor 9 attB mouse shown in upper right corner of F. Quantification of human Factor 9 positive area in livers of 3 treated mice shown in G.

In summary, here we describe a previously unreported limitation using RTs to write large (~40bp) sequences in non-dividing mouse hepatocytes. It is currently not clear why this issue is specific to rodent hepatocytes, as high beacon fidelity was achieved in both human and cyno primary hepatocytes. We speculate that this is likely due to host cell DNA repair machinery, or the low levels of dNTPs in non-dividing cells. In the context of PGI, we were able to overcome this issue either by dosing neonatal mice (cycling cells), or by engineering mice with pre-installed beacons. Integrase-mediated gene insertion was efficient in these models and was able to result in the expression of integrated transgene levels by harnessing endogenous promoters. An additional method to overcome this limitation is to use an alternate editing system such as a ligase-mediated editing [16]. We have identified a previously unreported challenge in the field and provided solutions which can pave the path for efficient translation and development of large genomic integration.

## Materials and Methods

### mRNA synthesis and purification

In-vitro transcription (IVT) was performed to synthesize mRNA from template plasmid DNA (GenScript) that had been linearized using the BspQI enzyme. Following synthesis, mRNA was purified by tangential flow filtration (TFF), followed by affinity chromatography using POROSTM Oligo (dT)25 Affinity Resin (Thermo Scientific). Ultrafiltration/diafiltration (UF/DF) was then carried out to concentrate the mRNA the 2 mg/mL and to exchange the mRNA into UltraPureTM DNase/RNase-Free Distilled Water (Invitrogen).

### LNP Formulation

Lipid nanoparticles were formulated using the NanoAssemblr Ignite system. Nucleic acid payloads were diluted in a 50 mM pH 4.5 acetate buffer. All lipids were purchased from commercial vendors: ALC-0315 (Broadpharm), DSPC (NOF America), cholesterol (Avanti), and DMG-PEG2000 (NOF America). Lipids were diluted in ethanol and combined to create a final stock solution according to the desired molar ratio. Mixing of lipids and nucleic acids was performed to achieve a final LNP composition with an N/P ratio=6. LNP were then diluted 1:1 with PBS and dialyzed overnight. LNP were concentrated using ultracentrifugation, diluted to the desired concentration using Tris-sucrose buffer pH 8.0, and sterile filtered prior to freezing. All LNP were analyzed using DLS and Ribogreen to assess size, polydispersity, and encapsulation efficiency of cargo and determined to meet the desired QC parameters.

### Cycling cell-based assays

HEK293 and Hepa1-6 were purchased from ATCC and cultured in DMEM (Gibco) with 10% fetal bovine serum (Gibco One Shot).

### Primary cell-based assays

Cryopreserved primary mouse hepatocytes (MSCP10 MC945, Gibco) were recovered in Hepatocyte Plating Media (A1217601, CM300, Gibco) and plated onto Collagen I coated 96-well tissue culture plates (356698, Corning) at 20k cells per well. Cryopreserved primary cynomolgus monkey hepatocytes (MKCP10 CY427, Gibco) were recovered in Hepatocyte Plating Media (A1217601, CM300, Gibco) and plated onto Collagen I coated 96-well tissue culture plates (356698, Corning) at 48k cells per well. Cryopreserved primary human hepatocytes (HMCPMS, Hu8450, Gibco) were recovered in Cryopreserved Hepatocyte

Recovery Media (CM7000, Gibco) and plated at 42k cells per well. 8 hours post recovery cells were washed and cultured in maintenance media (A1217601, CM400, A2737501, Gibco). Cells were treated with AAV at 1e6 Multiplicity of Infection 1 day after plating and transfected with nCas9-RT mRNA, atgRNA 1, atgRNA 2, and Bxb1 mRNA using

Lipofectamine Messenger MAX (LMRNA001, Invitrogen) in Opti-MEM (31985062, Gibco) 2 days after plating. Cells were collected for analysis 5 days after transfection.

### Animal studies

Transgenic C57BL/6J mice with a knock-in of the attB site in intron 1 of F9, Rosa26 or Pah were generated by Biocytogen (Beijing, China) using CRISPR/Cas9. Mice were transferred to Biomere (Worcester, MA, USA) for breeding. Mice at least 6 weeks old were injected intravenously via tail vein with AAV8 encoding the template DNA followed by administration of LNPs containing the integrase mRNA a week later. All animals underwent euthanasia 7 days after LNP administration and liver lobes were harvested for analysis.

Neonatal mice were injected intravenously into facial vein at 2 days old with LNP1 (nCas9-RT and guides) for beacon placement. At 6 weeks old, mice were injected intravenously via tail vein with AAV8 encoding the template DNA followed by administration of LNP2 containing the integrase mRNA a week later. All animals underwent euthanasia 7 days after LNP administration and liver lobes were harvested for analysis.

### gDNA isolation

Cells were lysed with QuickExtract (QE0905T, Lucigen) and gDNA was purified using SPRI magnetic bead cleanup.

Liver tissue was homogenized on Precellys Evolution (cat K002198-PEVO0-A.0 Combo, Bertin technologies, WA, USA). DNA was extracted with quick-DNA/RNA MagBead kit (cat R2131, Zymo research, CA, USA).

### Droplet Digital Polymerase Chain Reaction (ddPCR) analysis

Custom primers and probes were designed to measure editing. Results were normalized to custom reference assays targeting unedited regions of the same genes in the respective species. Probes were dual labelled with 3′-3IABkFQ and either 5′-carboxyfluorescein (FAM) for edit targets or 5′-hexachloro-fluorescein phosphoramidite (HEX) for reference. Assays were validated using gBlocks representing edit outcomes to test for both specificity and linearity. All primers, probes, and gBlocks were synthesized by IDT (Coralville, IA, USA). The reaction mix for all reactions, was composed of 12 µL of 2x ddPCR Supermix for probes (No dUTP) (cat 1863025 Bio-Rad, Hercules, CA, USA), 1.2 µL of each primer and probe mix, 0.12 µL of HindIII (cat FD0505, Thermo Fisher Scientific, MA, USA), 0.12 µL Eco91I (cat FD0394, Thermo Fisher Scientific, MA, USA), 1 µL genomic DNA and water to a final volume of 25 µL. Droplets were generated on the AutoDG Instrument for automated droplet generation (cat 186410, Bio-Rad, Hercules, CA, USA). PCR amplification was performed with the following initial denaturation at 95 °C for 10 min, followed by 40 cycles of denaturation at 94 °C for 30 s and combined annealing/extension step at 58 °C for 1 min, and a final step at 98 °C for 10 min, ending at 4 °C.

### HiBiT quantification

HiBiT from integrated PAH cargo was detected using the Nano-Glo HiBiT lytic detection system (N3050, Promega, Madison, WI, USA) following manufacturer protocol. For cell-based assays, 50 μL of master mix containing 100:2:1 ratio of Nano-Glo HibiT Lytic Buffer, Substrate, and LgBiT Protein was added per cell well and incubated for 10 minutes while shaking and read after 10 minutes additional incubation using GloMax Explorer (cat GM3500, Promega, Madison, WI, USA) with 0.3s integration. For in vivo studies, 125 μL of master mix was used for 25 µL of serum with all other steps identical.

### Human F9 ELISA

Human F9 in mouse serum was quantified by ELISA using hF9 ELISA kit (cat LS-F22871, Lifespan Biosciences, Lynnwood, WA, USA) following the manufacturer’s protocol. Mouse serum was diluted 10-fold in Reference Standard & Sample Diluent, provided by the kit.

Expression of hF9 was normalized to human serum.

### Immunofluorescence Imaging

Immunofluorescence to detect hF9 in mouse liver was developed by Invicro (Needham, MA, USA). Briefly, mouse left liver lobe was fixed in 10% neutral buffered formalin solution for 24 hours. Tissues were washed and stored in 70% ethanol. Tissues were sectioned and mounted on microscope slides. Sections were permeabilized in 0.1% Triton X-100 in PBS. Primary antibody Anti-human Factor IX (GAFIX-AP, Affinity Biologicals Inc, Ancaster, ON Canada) at 1:5000 was added on slides. Slides were washed. Secondary anti-Goat HRP Ab (DISCOVERY OmniMap anti-Gt HRP, Ventana Medical Systems, Oro Valley, AZ, USA) was added. Slides were washed again and Hoechst 1:500 in water was added for nucleus detection. Slides were visualized under microscope. Quantification of the hF9 was performed by calculating the percent hF9 positive area over total area.

## References

1. Gao C: Genome editing in crops: from bench to field. National Science Review 2014, 2(1):13–15.

2. Badwal AK, Singh S: A comprehensive review on the current status of CRISPR based clinical trials for rare diseases. Int J Biol Macromol 2024, 277(Pt 2):134097.

3. Xu L, Yao S, Ding YE, Xie M, Feng D, Sha P, Tan L, Bei F, Yao Y: Designing and optimizing AAV-mediated gene therapy for neurodegenerative diseases: from bench to bedside. J Transl Med 2024, 22(1):866.

4. Maeder ML, Stefanidakis M, Wilson CJ, Baral R, Barrera LA, Bounoutas GS, Bumcrot D, Chao H, Ciulla DM, DaSilva JA et al: Development of a gene-editing approach to restore vision loss in Leber congenital amaurosis type 10. Nat Med 2019, 25(2):229–233.

5. Metais JY, Doerfler PA, Mayuranathan T, Bauer DE, Fowler SC, Hsieh MM, Katta V, Keriwala S, Lazzarotto CR, Luk K et al: Genome editing of HBG1 and HBG2 to induce fetal hemoglobin. Blood Adv 2019, 3(21):3379–3392.

6. Yarnall MTN, Ioannidi EI, Schmitt-Ulms C, Krajeski RN, Lim J, Villiger L, Zhou W, Jiang K, Garushyants SK, Roberts N et al: Drag-and-drop genome insertion of large sequences without double-strand DNA cleavage using CRISPR-directed integrases. Nat Biotechnol 2023, 41(4):500–512.

7. Pandey S, Gao XD, Krasnow NA, McElroy A, Tao YA, Duby JE, Steinbeck BJ, McCreary J, Pierce SE, Tolar J et al: Efficient site-specific integration of large genes in mammalian cells via continuously evolved recombinases and prime editing. Nat Biomed Eng 2024.

8. Jenny Xie, Maike Thamsen Dunyak, Patrick Hanna, Angela X. Nan, Brett Estes, Jesse C. Cochrane, Shuai Wu, Jie Wang, Connor McGinnis, Qiang Wang, Rejina Pokharel, Dev Paudel, Jason Zhang, Dan Li, Parth Amin, Siddharth Narayan, Angela Hsia, Dane Z. Hazelbaker, Xiarong Shi, Meredith Packer, Brian Duke, Ryan Dickerson, Charlotte Piard, Martin Meagher, Jason Gatlin, Sonke Svenson, Adrianne Monsef, Raymond W. Bourdeau, Kieu Lam, Steve Reid, Mohammad Kazemian, Nisher Chander, Richard Holland, James Heyes, Swati Mukherjee, Sandeep Kumar, Daniel J. O’Connell, Jonathan D. Finn Curative levels of endogenous gene replacement achieved in non-human primate liver using programmable genomic integration. bioRxiv 2024, 10.1101/2024.10.12.617700.

9. Bock D, Rothgangl T, Villiger L, Schmidheini L, Matsushita M, Mathis N, Ioannidi E, Rimann N, Grisch-Chan HM, Kreutzer S et al: In vivo prime editing of a metabolic liver disease in mice. Sci Transl Med 2022, 14(636):eabl9238.

10. Davis JR, Banskota S, Levy JM, Newby GA, Wang X, Anzalone AV, Nelson AT, Chen PJ, Hennes AD, An M et al: Efficient prime editing in mouse brain, liver and heart with dual AAVs. Nat Biotechnol 2024, 42(2):253–264.

11. MacRae SL, Croken MM, Calder RB, Aliper A, Milholland B, White RR, Zhavoronkov A, Gladyshev VN, Seluanov A, Gorbunova V et al: DNA repair in species with extreme lifespan differences. Aging (Albany NY) 2015, 7(12):1171–1184.

12. Roesch F, Schwartz O: The SAMHD1 knockout mouse model: in vivo veritas? EMBO J 2013, 32(18):2427–2429.

13. Miazzi C, Ferraro P, Pontarin G, Rampazzo C, Reichard P, Bianchi V: Allosteric regulation of the human and mouse deoxyribonucleotide triphosphohydrolase sterile alpha-motif/histidine-aspartate domain-containing protein 1 (SAMHD1). J Biol Chem 2014, 289(26):18339–18346.

14. Apte U, Zeng G, Thompson MD, Muller P, Micsenyi A, Cieply B, Kaestner KH, Monga SP: beta-Catenin is critical for early postnatal liver growth. Am J Physiol Gastrointest Liver Physiol 2007, 292(6):G1578–1585.

15. Wang MJ, Chen F, Lau JTY, Hu YP: Hepatocyte polyploidization and its association with pathophysiological processes. Cell Death Dis 2017, 8(5):e2805.

16. Angela Xinyi Nan MC, Christopher Bartolome, Neeta Shadija, Dan Li, Brett Estes, Jessica Von Stetina, Wei Li, Jason Andresen, Jesse C Cochrane, Chen Bai, Jason Gatlin, Jie Wang, Davood Norouzi, Sandeep Kumar, Maike Thamsen Dunyak, Leonard Chavez, Anmol Seth, Shakked Halperin, Jonathan D Finn, Jenny Xie: Ligase-mediated programmable genomic integration (L-PGI): an efficient site-specific gene editing system that overcomes the limitations of reverse transcriptase-based editing systems bioRxiv 2024, 10.1101/2024.09.27.615478.

