## Supplemental figure 1 for "Low RT-based Genome Editing Fidelity in Mouse Hepatocytes: Challenges and Solutions"

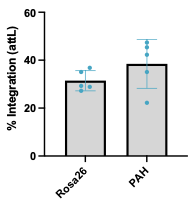

Supplemental Fig 1. I-PGI in transgenic attB beacon mice. Integration efficiency in transgenic mice with intact attB site placed in Intron 1 of Rosa26 and Pah. Mice were treated with AAV containing the template DNA (2e13 vg/kg). Human Factor 9 encoding AAV was used for Rosa26 attB mice and human Pah encoding AAV was used for Pah AttB mice. 1 week later mice were dosed with 3mg/kg LNPs containing Integrase mRNA. Integration efficiency was assessed in liver 1 week post LNP dosing.
